# Ledidi: Designing genomic edits that induce functional activity

**DOI:** 10.1101/2020.05.21.109686

**Authors:** Jacob Schreiber, Yang Young Lu, William Stafford Noble

## Abstract

The development of modern genome editing tools has enabled researchers to make edits with high precision, but has left unsolved the problem of designing these edits. We propose Ledidi, an approach that treats the design of genomic edits as an optimization problem where the goal is to produce the desired output from a predictive model. The discrete nature of biological sequences makes direct optimization challenging, but we overcome this by using the Gumbel-Softmax reparameterization trick. We validate Ledidi by pairing it with the Basenji model, which makes predictions for thousands of functional profiles, and designing edits that affect CTCF binding and induce cell type-specific binding of JUND.

## 1. Introduction

Recent advances in genome editing, driven by the discovery and development of the CRISPR-Cas9 system, have significantly reduced the cost of precisely modifying genomic sequence within living cells (Adli, 2018). Consequently, genome editing is now routinely used in a variety of contexts, including engineering crops (Arora & Narula, 2017), development of theraupeutics (Behan et al., 2019; Szlachta et al., 2018), and scientific investigation (Hsu et al., 2014). One such area of scientific investigation involves linking sequence elements to functional activity such as protein binding, chromatin accessibility, and transcription.

Although scientists use genome editing methods to great effect, these methods face several challenges. The first is the difficulty in knowing precisely what edits will yield a desired effect. A second challenge is that the editing process sometimes produces unintended consequences. For instance, deleting a protein binding site will likely affect protein binding, but it may also inadvertantly influence chromatin accessibility or transcription. Even small changes in the sequence of the genome can cause major effects.

Understanding the effects that small modifications, even single-nucleotide variants, have on functional profiles has been phrased as a supervised machine learning task. In this formulation, the input to a model is a window of nucleotide sequence and the output is the signal from a panel of functional assays. A recent model, Bassenji (Kelley et al., 2018), treats the problem as a regression task by predicting signal across > 131 kb of sequence at a resolution of 128 bp. These models can quickly predict the effect of mutations and other genomic alterations on biochemical activity in a given region, but require searching through an exponential number of combinations to find the best set of edits that yield a desired outcome.

Several approaches have since been proposed for quickly designing biological sequences that exhibit desired characteristics. These methods generally consist of two components: a generation step that produces sequences and an *oracle*, i.e. a model that can quickly evaluate the quality of each proposed sequence. One such method, deep exploration networks (Linder et al.), involves a pair of networks that generate and evaluate sequences in a fashion similar to that of a generative adversarial network. The method achieves diversity by penalizing the generation of similar pairs of sequences. An alternate approach relaxes the requirement that the oracle be differentiable through the use of an adaptive sampling method that is based on parametric conditional density estimation (Brookes et al., 2019). A third approach, the encoder-decoder-analyzer model (Gupta & Kundaje, 2019), involves the training of three neural networks to encode the sequence, decode the sequence, and perturb the internal latent state such that the resulting sequence exhibits the desired characteristics.

We propose a method, named Ledidi, for designing edits to the genome that induce a desired functional landscape. Ledidi phrases the design task as an optimization problem where the goal is to identify a set of edits that results in the desired functional profiles. Because experimental validation of each proposed sequence is infeasible, Ledidi relies on an oracle. We chose Basenji as the oracle model because it provides a detailed output and thus fine-grained design over the design process. A difficulty in this optimization problem is that genomic sequence is discrete; however, we overcome this difficulty by using the Gumbel-Softmax reparameterization trick (Jang et al., 2016) that enables standard gradient descent methods to be used on discrete inputs. An important distinction between Ledidi and the works of Linder et al. and Brookes et al. (2019) is that Ledidi does not design entire sequences but rather a small set of edits, and in contrast to the work of Gupta & Kundaje (2019), Ledidi does not involve training a new model.

## 2. Methods

Ledidi aims to design a small set of edits that change the output of a pre-trained model in a desired manner. Thus, Ledidi optimizes an objective function that is a mixture of two terms: the first term is the distance between the original sequence and the proposed edited sequence (the sequence loss), and the second term is the distance between the model output using the edited sequence and the desired output (the functional loss). More formally, the original sequence is *X*_0_ ∈ {0,1}^*n×d*^, where *n* is the length of the sequence and *d* is the size of the alphabet; the edited sequence is *X* ∈ {0, 1}^*n×d*^; the model output using the edited sequence is 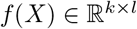, where *k* is the size of the output sequence and *l* is the number of output measurements per position, and the desired output 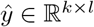. The objective function for this optimization is

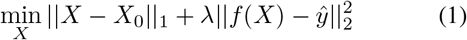

The second term is weighted by λ to either encourage a small number of edits when small or a closer match to the desired output when large. Unfortunately, Equation 1 is difficult to optimize directly because *X* is discrete.

We overcome this challenge by using a relaxation of the discrete representation of *X* that approximates the behavior of the one-hot encoding. This relaxation, known as the Gumbel-Softmax reparameterization trick (Jang et al., 2016), involves substituting a new, continuous various, 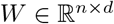, for *X*, where

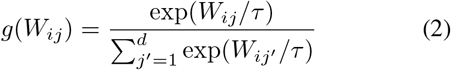

and *W* is initially set to *X*_0_. The function *g*(*W*) is parameterized by a positive temperature, *τ*, that controls how close to a discrete representation the Gumbel-Softmax distribution is. As *τ* approaches 0, *g*(*W_ij_*) smoothly approaches the discrete distribution {0,1} and as *τ* approaches infinity, *g*(*W_ij_*) approaches the uniform distribution.

A discrete sequence *X* can be recovered from *g*(*W*) using a simple argmax operation.

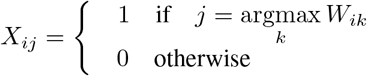

With this new variable *W*, we can solve an alternative form of Equation 1:

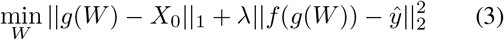

Equation 3 can be solved by standard gradient descent methods. In this work, we use Nesterov’s accelerated gradient descent (Nesterov, 1983) with a maximum number of iterations of 100 and a patience of 10.

While Ledidi is general purpose enough to work with any model and sequence size, we chose to use Basenji (Kelley et al., 2018). Basenji takes *X*_0_ ∈ {0, 1}^131,072×4^ as input, performs deep neural network operations *f*, and predicts the functional profile 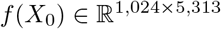.

In our experiments we focused on designing edits that affect only a subset of the functional profiles predicted by Basenji. Accordingly, we define a mask *M* ∈ {0, 1}^1,024×5,313^ that is applied elementwise to the loss, giving the final objective

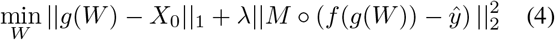

During the optimization process differences arise between the loss calculated using *g*(*W*) and *X* as input to the model: specifically, *g*(*W*) may yield more accurate functional profiles than the corresponding discretized sequence *X* because *g*(*W*) is directly optimized. Indeed, we found that frequently this difference resulted in the best *X* being found midway through the optimization process. We mitigate this issue by optimizing the objective function given in Equation 4 normally but returning the *X* that gave the smallest loss at any point.

## 3. Results

### 3.1. CTCF binding can be deleted and induced

Our first evaluation of Ledidi involved designing edits that delete or induce binding of the CTCF protein. CTCF has two properties that make it an ideal candidate for an initial evaluation: first, the protein binds tightly to a known motif, and second, this binding generally occurs at similar loci across cell types. These two factors indicate a strong connection between genomic sequence and functional activity. For this evaluation, we identified 53 high confidence (q-value < 0.01) CTCF motif sites across the human genome (build hg38) using the motif scanning tool FIMO (Grant et al., 2011). We obtained Basenji’s predictions by running ~ 131kb sequences centered around these 53 sites to get predictions of CTCF binding in the cell line GM12878. Next, we constructed our desired CTCF profiles for knock-out experiments by zeroing out the predicted fold-change values for CTCF from Basenji in a window of ±25 positions surrounding the motif (Fig 1A), and for knock-in experiments by copying the peak in that window to 6.4 kb upstream.

**Figure 1.**
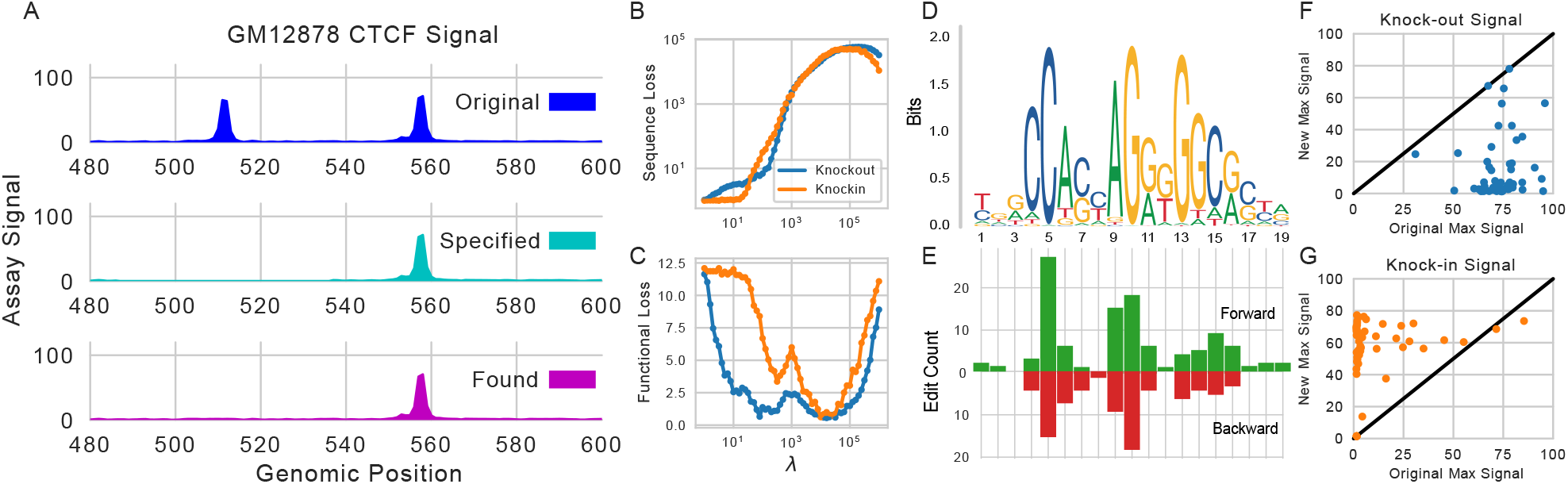
Editing CTCF motif sites. (A) Ledidi takes as input the initial sequence (not shown) and the desired signal. The output is the edited sequence (not shown) and the predicted signal for that sequence. (B) The sequence loss (the first term in Equation 1) averaged over all 53 loci for different values of λ on CTCF knock-outs and knock-ins. (C) Similar to B, except displaying the functional loss (the second term in Equation 1). (D) The CTCF motif and (E) the distribution of edits proposed by Ledidi when knocking out CTCF peaks, corrected for the motif occuring on the forward or reverse strands. (F) The original signal of each of the 53 regions and the new signal arising from Ledidi’s edits when knocking out a peak. (G) Similar to F except for inducing a peak.

We began by quantifying the effect that changing Ledidi’s mixing parameter λ has on the proposed set of edits. We ran Ledidi using the 53 extracted sequences and desired knock-out/knock-in CTCF profiles using the default parameters of *τ* = 0.05 and a learning rate of 10^−3^ but varied λ from 1 to 10^6^. When we plotted the average sequence loss (i.e., the number of edits) and functional loss of the returned sequences we observed that increasing λ led to a larger set of proposed edits (Fig 1B) and a W-shaped curve in the functional loss (Fig 1C) for both tasks. The smaller number of edits made in the first dip suggests that these sequences were designed by making a small number of targeted changes, whereas the second dip corresponds to the trivial solution of redesigning large portions of the input sequence. Potentially, the increase in loss between the two dips corresponds to the boundary between these two regimes.

Guided by the results of the hyperparameter sweep, we focused on the knock-outs designed when using a λ of 100. On average, Ledidi proposed 5.85 edits per locus, with 190 of the 310 edits occuring within the CTCF motif that the window was centered on. These edits, which occured primarily on the most conserved nucleotides in the CTCF motif on both strands (Fig 1D/E), reduced the maximum signal within the window by an average fold change of 58.3, and all except for 2 loci exhibited diminished signal (Fig 1G). During optimization, we noted that the functional loss calculated using *g*(*W*) would steadily decrease for both of these loci but that the loss calculated using *X* did not.

Next, we examined the edits designed to induce CTCF binding (Fig 1G) when using a λ of 333, a setting that exhibited good performance in the hyperparameter sweep. Encouragingly, these edits increased the maximum CTCF signal at the target by an average fold change of 44.5, and only 7 loci did not exhibit increased signal. At one locus, we observed that the proposed edits decreased the maximum signal from 85.4 to 73.6; however, because the desired signal had a maximum signal of 77.7, these edits actually resulted in a better sequence despite the decrease in signal.

### 3.2. JUND binding can be deleted in a biosample-specific manner

Although our CTCF experiments served as a useful initial validation of Ledidi, a more compelling use case involved editing the genome to produce complex functional profiles. Thus, we next turned to the more challenging task of inducing cell type-specific protein binding. The protein JUND is a good target for this task because JUND has been shown to recognize different sequence motifs in different cell types (Arvey et al., 2012), potentially due to interactions with different sets of co-factors.

In this evaluation, we used Ledidi to knock out the binding of JUND in h1-hESC while preserving its binding in GM12878 at sites where JUND normally binds in both cell types. We extracted the sequence surrounding 23 sites, one from each chromosome, where each site was the location of the strongest corresponding ChIP-seq signal p-value signal added over both cell types for JUND. Accordingly, we zeroed-out the central peak for h1-hESC but retained the peak in GM12878 (Fig 2A).

Intriguingly, we found that Ledidi was capable of performing this knock-out with relative ease, and that there were cases where nearby peaks that extended into the zeroed out region were preserved while the punctate peaks within the zeroed out region were removed (Fig 2B). This finding suggested that Ledidi is not simply finding an out-of-distribution sequence pattern that results in a low signal output. When we comprehensively applied Ledidi to the 23 loci, we found that the resulting edits reduced the signal within the zeroed out regions in h1-hESC quite well, with much smaller impacts to those regions in GM12878. Notably, we had to change the hyperparameters of Ledidi: we set the learning rate to 10^−6^, the maximum number of iterations to 1,000, and λ to 10^5^. In contrast to eliminating CTCF binding, which took only a few edits per locus, designing edits that induce biosample specific binding of JUND required between 78 and 670 edits per locus, indicating that it was a more challenging task. Encouragingly, these edits occured primarily in the region that was being perturbed (Fig 2C).

**Figure 2.**
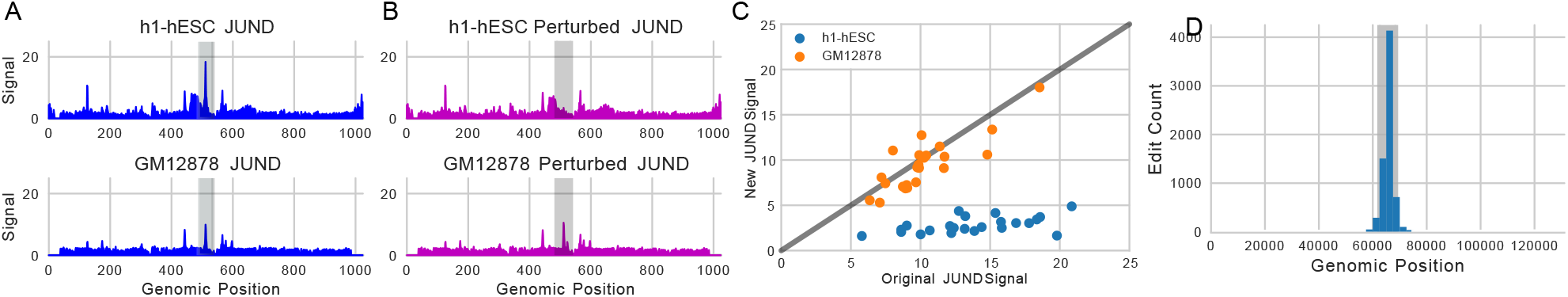
Editing JUND binding in two cell types. (A) An example of the original JUND signal predicted by the Basenji model. A shaded grey box indicates the region that is zeroed out in h1-hESC. (B) The resulting predicted signal after the sequence is edited by Ledidi, with the same region shaded. (C) The maximum signal value in the region that was zeroed out in h1-hESC for both h1-hESC and GM12878, showing the change in signal after the sequence edits. (D) A histogram over the relative loci of the input to Basenji where the edits are made. The shaded region shows where the signal was zeroed out.

## 4. Discussion

In this work we propose Ledidi, a method for designing edits to a discrete input sequence that result in a desired change in model output. We first demonstrate that Ledidi works by successfully designing genomic edits that induce and delete CTCF binding. Then, we successfully apply Ledidi to the more challenging task of inducing cell type-specific binding of JUND, a protein that is known to bind at different loci in different cell types. These examples provide empirical evidence for the usefulness of Ledidi.

Although Ledidi generally achieved reasonable results, we observed that the hyperparameters had to be changed as the tasks became more complex. Naturally, we found that increasing λ with the number of edits anticipated for successful design was important. We also found that decreasing the learning rate could lead to more accurate outputs that also required fewer edits, but had the drawback of (sometimes dramatically) increasing runtime. We found that a good strategy was to initially begin with a large λ and then decrease both the learning rate and λ in successive optimization runs to find a compact set of edits. Finding ways of automatically tuning these hyperparameters, such as through annealing schedules, will likely be important for out-of-the-box use of Ledidi by non-specialists.

Ledidi has a few limitations. First, Ledidi cannot currently propose sequence insertions or deletions. Second, any deficiencies in the oracle that Ledidi uses will propogate into the proposed edits. For instance, we observed that the Basenji model did not predict a significant difference between the binding of MYC and MAX, despite subtle but known differences in their binding motifs (Allevato et al., 2017). Finally, the speed of Ledidi is limited by the inference step of the oracle. The same complexity that made Basenji an accurate model also makes the inference step longer. Using a GTX 2080 Ti GPU, the design process took up to a minute for simple deletions and several minutes for more complex edits. Fortunately, the second and third limitations can both be overcome by using a small model tuned for the relevant phenomena.

It is worth mentioning that Ledidi relies on the generalization capabilities of the oracle that is being used. When the oracle generalizes well, edited sequences can be easily validated and those sequences that do not yield the desired change, potentially because of an issue during optimization or because the desired change is impossible, can be flagged. However, when the oracle does not generalize well, Ledidi is likely to propose edits that do not, in reality, induce the desired activity. Therefore, an important area of future work is detecting when Ledidi is proposing edits that are out-of-distribution (Bulusu et al., 2020) for the oracle that is being used.

We have made our code available at https://github.com/jmschrei/ledidi.

## References

Adli, M. The CRISPR tool kit for genome editing and beyond. Nature Communications, 9(1911), 2018.

Allevato, M., Bolotin, E., Grossman, M., Mane-Padros, D., Sladek, F M., and Martinez, E. Sequence-specific DNA binding by MYC/MAX to low-affinity non-e-box motifs. PLoS One, 12(7):e0180147, 2017.

Arora, L. and Narula, A. Gene editing and crop improvement using CRISPR-Cas9 system. Front Plant Sci., 8 (1932), 2017.

Arvey, A., Agius, P., Noble, W. S., and Leslie, C. Sequence and chromatin determinants of cell-type specific transcription factor binding. Genome Research, 22(9):1723–1734, 2012.

Behan, F. M., Iorio, F., and et al., G. P. Prioritization of cancer therapeutic targets using CRISPR-Cas9 screens. Nature, 568:511–516, 2019.

Brookes, D. H., Park, H., and Listgarten, J. Conditioning by adaptive sampling for robust design. In Proceedings of the 36th International Conference on Machine Learning, volume 97, pp. 773–782, 2019.

Bulusu, S., Kailkhura, B., Li, B., Varshney, P. K., and Song, D. Anomalous instance detection in deep learning: a survey. arXiv, 2020.

Grant, C. E., Bailey, T. L., and Noble, W. S. FIMO: Scanning for occurrences of a given motif. Bioinformatics, 27 (7):1017–1018, 2011.

Gupta, A. and Kundaje, A. Targeted optimization of regulatory DNA sequences with neural editing architectures. biorXiv, 2019.

Hsu, P. D., Lander, E. S., and Zhang, F. Development and applications of CRISPR-Cas9 for genome engineering. Cell, 157(6):1262–1278, 2014.

Jang, E., Gu, S., and Poole, B. Categorical reparameterization with Gumbel-Softmax. arXiv, 2016. arXiv:1611.01144.

Kelley, D., Reshef, Y., Bileschi, M., Belanger, D., McLean, C., and Snoek, J. Sequential regulatory activity prediction across chromosomes with convolutional neural networks. Genome Research, 2018.

Linder, J., Bogard, N., Rosenberg, A. B., and Seelig, G. Deep exploration networks for rapid engineering of functional DNA sequences. bioRxiv. https://www.biorxiv.org/content/10.1101/864363v1.

Nesterov, Y. A method for solving a convex programming problem with convergence rate o(1/k^2^). Soviet Mathematics Doklady, 27:372–376, 1983.

Szlachta, K., Kuscu, C., and et al., T. T. CRISPR knockout screening identifies combinatorial drug targets in pancreatic cancer and models cellular drug response. Nature Communications, 9(4275), 2018.

